# 3D printed lung on a chip device with a stretchable nanofibrous membrane for modeling ventilator induced lung injury

**DOI:** 10.1101/2021.07.02.450873

**Authors:** Sinem Tas, Emil Rehnberg, Deniz A. Bölükbas, Jason P. Beech, Liora Nasi Kazado, Isak Svenningsson, Martin Arvidsson, Axel Sandberg, Kajsa A. Dahlgren, Alexander Edthofer, Anna Gustafsson, Hanna Isaksson, Jeffery A. Wood, Jonas O. Tegenfeldt, Darcy E. Wagner

**Affiliations:** Lung Bioengineering and Regeneration, Department of Experimental Medical Sciences, Stem Cell Centre, Wallenberg Center for Molecular Medicine, Lund University, Lund, 22362 Sweden; Division of Solid State Physics and NanoLund, Physics Department, Lund University, PO Box 118, 22100 Lund, Sweden; Department of Biomedical Engineering, Lund University, Box 118, 221 00 Lund, Sweden; Soft Matter, Fluidics and Interfaces, MESA+ Institute for Nanotechnology, University of Twente, Enschede, 7522 NB, The Netherlands

## Abstract

Mechanical ventilation is often required in patients with pulmonary disease to maintain adequate gas exchange. Despite improved knowledge regarding the risks of over ventilating the lung, ventilator induced lung injury (VILI) remains a major clinical problem due to inhomogeneities within the diseased lung itself as well as the need to increase pressure or volume of oxygen to the lung as a life-saving measure. VILI is characterized by increased physical forces exerted within the lung, which results in cell death, inflammation and long-term fibrotic remodeling. Animal models can be used to study VILI, but it is challenging to distinguish the contributions of individual cell types in such a setup. *In vitro* models, which allow for controlled stretching of specific lung cell types have emerged as a potential option, but these models and the membranes used in them are unable to recapitulate some key features of the lung such as the 3D nanofibrous structure of the alveolar basement membrane while also allowing for cells to be cultured at an air liquid interface (ALI) and undergo increased mechanical stretch that mimics VILI. Here we develop a lung on a chip device with a nanofibrous synthetic membrane to provide ALI conditions and controllable stretching, including injurious stretching mimicking VILI. The lung on a chip device consists of a thin (i.e. ∼20 µm) stretchable poly(caprolactone) (PCL) nanofibrous membrane placed between two channels fabricated in polydimethylsiloxane (PDMS) using 3D printed molds. We demonstrate that this lung on a chip device can be used to induce mechanotrauma in lung epithelial cells due to cyclic pathophysiologic stretch (∼25%) that mimics clinical VILI. Pathophysiologic stretch induces cell injury and subsequently cell death, which results in loss of the epithelial monolayer, a feature mimicking the early stages of VILI. We also validate the potential of our lung on a chip device to be used to explore cellular pathways known to be altered with mechanical stretch and show that pathophysiologic stretch of lung epithelial cells causes nuclear translocation of the mechanotransducers YAP/TAZ. In conclusion, we show that a breathable lung on a chip device with a nanofibrous membrane can be easily fabricated using 3D printing of the lung on a chip molds and that this model can be used to explore pathomechanisms in mechanically induced lung injury.

## Introduction

The lung is a mechanically dynamic organ and subjected to several physical forces including mechanical stretch from breathing, shear stress from pulmonary blood flow, and surface tension due to alveolar lining fluid. Mechanical stretch within a physiological range (5-10%) is known to play a pivotal role in lung tissue development as well as in maintaining tissue homeostasis and directing adult regeneration.^1^ However, mechanical stretch which exceeds the physiological range (> 20%) is known to significantly affect the lung at both the tissue and cellular level.^1^ This pathophysiological stretch of the lung is a major clinical problem and can occur due to the use of mechanical ventilation in patients with pulmonary dysfunction due to injury or disease. While mechanical ventilation is lifesaving in these patients, overdistension of the lung can occur due to the use of excessive volume or pressures as well as due to disease which can cause inhomogeneous distribution of airflow. Increased physical forces exerted on the lung as a consequence of mechanical ventilation can exacerbate pre-existing lung damage and cause further damage resulting in ventilator induced lung injury (VILI).^2^ VILI is characterized by an increase in tissue level inflammation during the initial acute phase of injury and in many patients is followed by a subsequent fibrotic response which can lead to irreversible lung damage.^3^ Thus far there are no treatment options for patients who develop VILI and this can be partly attributed to a lack of suitable models in which to study the cellular and molecular events which occur during VILI onset and progression.

Due to similarities of chest anatomy and branching architecture with humans, large animal models have been frequently used to understand the pathophysiology of VILI. However, the availability of species-specific reagents to measure inflammatory mediators, receptors, or other proteins are major limitations of animal models, as well as their high cost and the specialized expertise required to run these experiments. Moreover, many VILI models are limited to short time points (a few hours) due to concerns regarding animal welfare making the study of VILI under conditions and timeframes which mimic human disease challenging.^4^ Thus, it is important to design physiologically relevant *in vitro* models to study the pathophysiology of VILI on a cellular level and which can be used to identify new treatments.

Several different *in vitro* lung tissue models with cyclic mechanical stretch have been developed by integrating manufacturing techniques with cell culture.^5-14^ Currently utilized models span from macro to microscale devices, the latter of which allow for complex devices to be manufactured with similar dimensions to the natural cell microenvironment. These devices have been used to study alveolar barrier function,^10, 11, 15^ lung disease^13, 16^, lung injury^7, 14^ and drug response^16, 17^ using lung on a chip devices. However, the study of pathological mechanical stretch on biologically relevant membranes has thus far been challenging in these setups due to the use of 2D planar membranes which do not mimic the nanofibrous basement membrane of the alveolus that can also undergo cyclic mechanical stretch within the ranges known to be present in VILI.

The majority of these previous devices, including lung on a chip devices, have used a thin, porous and stretchable polydimethylsiloxane (PDMS) membrane due to their good elastic, optical and biocompatibility properties.^1, 18^ However, due to their 2D planar surface, PDMS membranes do not accurately mimic the 3D natural ECM that cells grow on.^7, 10, 19 20^ These membranes typically lack the fibrous structure of the ECM and do not contain cell binding motifs. To remedy this, cell binding sites can be chemically incorporated onto surfaces (e.g. RGD) or ECM proteins can be incorporated directly during manufacture (e.g. blending) or post-manufacture through surface coating with ECM proteins (such as collagen, fibronectin) before cell seeding.^10, 12, 13^ In the latter approach, this thin ECM coating can help mimic native binding sites, but it does not mimic the 3D topography that cells experience *in vivo*. More recently, a thin hydrogel membrane made of ECM proteins of collagen and elastin has been developed with stretchable properties as a next generation membrane for use in a lung on a chip device to more closely mimic the alveolar air-blood barrier.^15^ While these membranes were stretchable up to 10%, the ECM hydrogel was fabricated using spontaneous thermal assembly of ECM molecules and thus lacked covalent crosslinking between ECM molecules, which rendered them somewhat mechanically weak and unable to mimic VILI where stretching exceeds 20%.^18, 21-23^

Therefore, materials which can satisfy the multiple critical aspects of the lung are needed; namely nanofibrous topography which supports cell growth while allowing for mechanical stretch in the range of that encountered during normal and pathologic scenarios. Electrospun membranes are one potential option due to the fact that they are made of micro-or nanofibers, where the electrospinning parameters and the material can be chosen to generate membranes with specific mechanical properties.^24^ This approach offers the further benefit that properties such as fiber diameter, fiber orientation, porosity, mechanical strength, stretchability and thickness can be tailored to mimic the properties of the native ECM.^25^ Electrospun membranes with fiber diameters in the micrometer range have previously been used to mimic the alveolar capillary basement membrane in macro-and microscale devices ^11, 26, 27^ However, only a few of these devices have incorporated basolateral media flow and these have been limited to macroscale devices.^28^ Such macroscale devices are costly and take up considerable space, and are therefore limited with regard to how many replicates can be run in parallel. This is a particularly important logistical aspect to consider for the use of primary cells in such devices which have limited availability and are known to change over passage number. Therefore, development of a next generation microscale (i.e. lung on a chip) device which has a nanofibrous membrane, which i) more closely mimics the topography of the native ECM, ii) allows for culture at air liquid interface and iii) can withstand normal and pathologic air and liquid shear stresses is needed in order to more closely mimic VILI *in vitro*.

Here, a lung on a chip device containing a stretchable nanofibrous membrane has been developed and investigated as a model of ventilator induced lung injury (VILI). The device consists of two parallel PDMS channels separated by a flexible, electrospun nanofibrous polycaprolactone (PCL) membrane for cell attachment. In order to reduce costs and eliminate the need for clean-room facilities, we used stereolithography based 3D printing to generate molds for PDMS casting. We show that epithelial cells can be grown at air-liquid interface, with an ability to control both air and media flow on respective sides of the membrane. For device validation, we exposed lung alveolar epithelial cells seeded on the air side of the chip to mechanical stretch comparable to what causes VILI (i.e. approximately 25%) and evaluated whether our lung on a chip could be used to induce pathologic features relevant to VILI. We found that pathologic stretch induced cell death and was able to induce nuclear translocation of the mechanotransducers YAP/TAZ. Therefore, the integration of a nanofibrous PCL membrane in a lung on a chip device, is a promising new approach for modeling VILI and could be used to further understand pathomechanisms and evaluate potential new therapies.

## Materials and Methods

### Lung on a chip device fabrication

The microfluidic device consisted of an electrospun membrane, which was sandwiched between two PDMS channels (**Figure 1a**). The molds for PDMS casting of the upper and lower chamber of the device were designed in the open access software Blender (**SI Figure1**) and printed from standard black resin (Formlabs) using a stereolithography 3D printer (Form 2, Formlabs). Next, the PDMS (Sylgard 184, Dow Corning) pre-polymer was mixed with its curing agent at a weight to weight ratio of 10:1 and degassed for at least 30 minutes, before pouring it into the mold. The PDMS was then cured for 1 hour at 80°C under normal atmospheric conditions in an oven. The fully cured PDMS was carefully peeled off from the 3D printed mold. A 1.5 mm hole puncher (World Precision Instruments) was used to generate holes to provide inlets and outlets of the upper and bottom channels.

**Figure 1:**
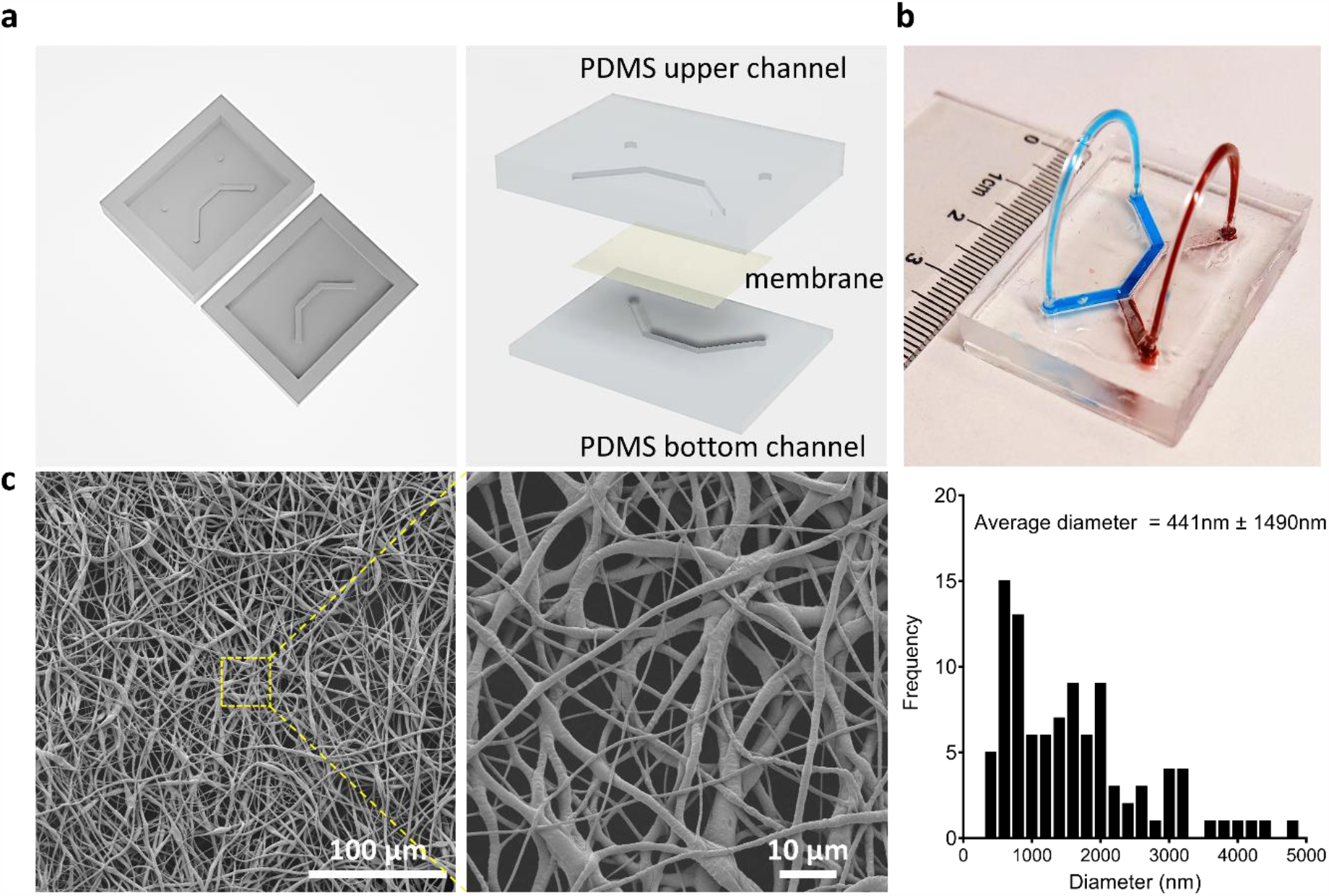
Microfluidic lung-on-a-chip device. **a)** 3D printed mold with a channel dimensions of 1.5 mm x 1mm x 10 mm (w x h x l). Two PDMS layers were aligned and irreversibly bonded to form two parallel channels separated by a ∼20 μm thick electrospun membrane. **b)** Representative image of an actual lung-on-a-chip microfluidic device. **c)** Representative SEM images of electrospun PCL membrane and fiber diameter distribution curve of electrospun PCL membranes (n=3).

In parallel, the same PDMS mixture (10:1) was spin coated on a glass slide at 800 rpm for 30s (WS-400-6NPP-LITE, Laurell Technologies Corporation) to cover the glass slide with a uniform coating of the PDMS mixture. The PDMS device parts were then placed on top of the PDMS covered glass slide to cover the surface of the thin layer of PDMS. The two parts were then placed inside an 80°C oven for 10-15 minutes, to partially cure the thin PDMS layer. A 20μm thick electrospun polycaprolactone (PCL) membrane (Cellevate AB, Sweden, 3D Nanomatrix™) was carefully placed over the channel on the top PDMS half of the chip. Once the membrane was put in place, the device was sealed by aligning the top and bottom channels. The assembled lung on a chip device was then left for 24 hours to fully cure inside of a ventilated space. Tygon tubing (ID: 0.5 mm and OD: 1.5 mm) was inserted into inlets and outlets of the device.

### Membrane morphology

The morphology and fiber diameters of the commercially available PCL membrane were evaluated using scanning electron microscopy (SEM). The PCL membranes were cut with dimensions of 50 mm x 50 mm. Changes in fiber orientation following mechanical stretch was assessed by applying stretch on to membranes assembled into the lung on a chip device (without cells) and stretched at 25 % and for 2 hours and compared to membranes assembled into the lung on a chip device but without stretch (n=3 for each condition). The samples were sputter-coated with 10 nm platinum/palladium (4:1 composition) (Quorum Q150T ES) and examined at an accelerating voltage of 3.0 kV using the secondary electron detector in a Jeol JSM-7800F FEG-SEM. The fibers were analyzed from images taken at 250× magnification using ImageJ to determine the mean fiber diameter (MFD) and the directionality plugin to generate fiber orientation distributions centered at 0°. A total of 90 bins, representing 2° increments, were used to generate histograms for each of the three independent experiments of membranes, stretched or unstretched.

### Mechanical testing

The mechanical properties of electrospun PCL membranes were determined using tensile testing (Instron 8511 load frame, High Wycombe, UK/MTS Test Star II controller, Minneapolis, US). A stack of 8 membranes was used for the tests, with a total thickness ranging between 80-100 μm (based on measured thickness with digital calipers). The PCL membranes (n=4) were cut into 20 mm wide strips with a gauge length of approximately 40 mm. The specimens were tested until failure at a speed of 10 mm/s and force-displacement curves were recorded. The stress-strain curves were calculated by normalizing the forces with the cross-sectional area and the displacement with the initial gauge length. The elastic modulus was calculated as the slope of the initial part of the stress-strain curve assuming an average sample thickness of 90 μm (thickness range 80-100 μm). The membranes displayed a bilinear mechanical response, and the yield strain was therefore determined from the intersection between two lines fitted to the elastic and plastic regions of the curve. The failure was taken as the maximum strain at failure.

### Numerical simulations

A numerical simulation was performed in order to estimate the membrane deflection in the lung on a chip device under applied pressure to model VILI. For simplicity, a 2D dimensional representative of the lung on a chip device was created in COMSOL Multiphysics 5.5™. The membrane geometry was constructed by using the actual chip dimensions as a width of 1.5 mm and length of 10 mm, while the height of the upper channel was taken as 1.0 mm. For simulations, the density (ρ), Young’s modulus (E), and Poisson ratio (ν) of the PCL membranes were set at ρ=1150 kg/m^3^, E=3 MPa and ν=0.442, respectively.^24, 29^ The deformation under applied pressure (gas) and flow was determined by solving a fluid-structure interaction problem. The PCL membrane was treated as a linear elastic material, while the fluid flow profile in the upper channel was determined through solution of the Navier-Stokes equations (in the laminar regime). The inlet flow to the channel/membrane region was considered as fully developed. Fluid-structure interaction was described by coupling fluid dynamics (Navier-Stokes equations) with structural mechanics (linear elastic material). The Navier-Stokes equations were solved using a P2-P1 basis (2^nd^ order Lagrange elements for velocity, 1^st^ order for pressure) while the deformation was solved using a 2^nd^ order Lagrange basis. The mesh consisted of 195 000 quadrilateral elements after successive refinement until the solution stopped changing.

Under the assumption that flow inside the microchannel is incompressible, steady, creeping and Newtonian, the Navier–Stokes equation which describes the flow behavior is given as follows,

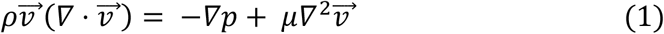

where ρ is the fluid density, 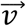 is the fluid velocity, p is the hydrodynamic pressure and μ is the fluid dynamic viscosity.

The membrane deformation was subject to the following boundary conditions:

1. no-slip boundary conditions applied on the rigid walls
2. zero displacement is prescribed along the PCL membrane-PDMS interface and the pressure is applied on top of the membrane to estimate the deflection
3. a fully developed velocity profile is specified

### Membrane stretching in the lung on a chip device

To confirm that the membrane could be mechanically stretched without causing delamination of the membrane from the two PDMS channels, the inlet of the top channel was connected to a syringe pump (World Precision Instruments) with a 2 mL syringe. The syringe pump was then set to deliver 150 μL at a respiratory rate of 6 breaths per minute (i.e. using a flow rate of 108 mL/h for dispensing followed by withdrawal of 150 μL at 108 mL/h). The membrane motion was then monitored using an inverted microscope (Nikon Eclipse Ts2R) using a 20x objective to confirm movement of the membrane. The chip was visually inspected for signs of air leakage or delamination of the top and bottom PDMS channels.

The displacement of the membrane was measured while the bottom channel was filled with liquid (i.e. water). The syringe pump was connected to the upper channel and 150 µL of air was injected at 108 mL/h and then subsequently withdrawn as described previously. Membrane movement was again monitored visually using an inverted microscope (Nikon Eclipse Ts2R) with a 20x objective and the chip was examined for signs of leakage.

### Measurement of membrane deflection

The membrane deflection was also experimentally determined. First, the outlet of the bottom channel was connected to Tygon tubing (ID: 0.5 mm and OD: 1.5 mm). The Tygon tubing connected to the bottom channel was then partially filled with red colored water to create a liquid plug. The colored water plug was then moved to approximately the center of the tubing by introducing air into the tubing. The inlet of the Tygon tubing was then connected to the outlet of the bottom channel and the outlet of the Tygon tubing (ID: 0.5 mm and OD: 1.5 mm) was left open to the atmosphere. The syringe pump was connected to the upper channel and 150 µL of air was injected at 108 mL/h and then subsequently withdrawn as described previously. The movement of the liquid plug was measured between every injection and withdrawal of air using millimeter graph paper. The displaced volume was then calculated using the inner diameter of the tubing to estimate membrane deflection.

Assuming the deflection of the membrane would be small and act as a linear elastic material, the deflection profile can be estimated to be parabolic and proportional to

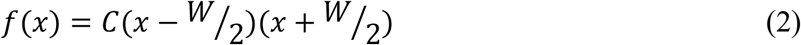

where *W* is the channel width. The constant *C* can be found from the displaced volume *V*.

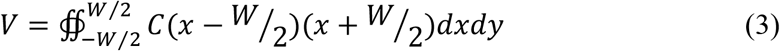

The deflection *h* is then given by

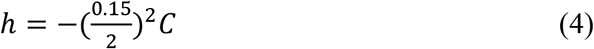

The length of the stretched membrane is then given by the parabolic arc length *s* which can be found from

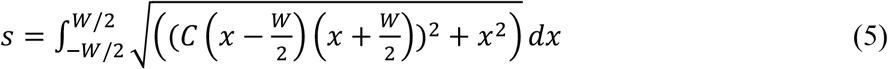

The membrane strain S_m_ can then be calculated from

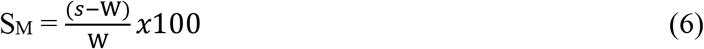

### Cell culture

Immortalized murine lung alveolar epithelial cells (MLE-12) (ATCC^®^ CRL-2110™) were used in all cell culture experiments. Cells were cultured in complete media comprised of DMEM/F12 (Gibco), 10% fetal bovine serum (FBS, Gibco) and 1% penicillin/streptomycin (P/S, Merck). MLE-12 cells were cultured until 90% confluence in tissue culture treated T75 flasks and then trypsinized, centrifuged, and resuspended in complete media prior to subsequent passaging and culture on electrospun membranes or cultured in the lung on a chip device.

In order to evaluate the ability of electrospun PCL membranes to be used as cell culture support, MLE-12 cells were seeded (3.0×10^5^ cells/cm^2^) onto electrospun PCL membrane inserts (Cellevate AB, Sweden, 3D Nanomatrix™) in 24 well cell culture plates (Sarstedt). Prior to cell seeding, membranes were sterilized with 70% ethanol for 30 minutes and washed twice with PBS. The membrane was incubated with complete cell culture media at 37°C for at least 30 minutes to adsorb proteins on the surface of the PCL membranes to support cell attachment. Cells were added to electrospun PCL membrane inserts in triplicates and cultured in an incubator (37°C, 5% CO_2_) for 72 hours to obtain a confluent monolayer. One media exchange was done 48 hours after the cell seeding.

### MTT metabolic activity assay

The MTT (thiazolyl blue tetrazolium bromide, Sigma Aldrich) assay was performed after 72 hours of incubation to analyze the metabolic activity of the cells and confirm cytocompatibility of the electropsun PCL scaffolds. 3.0×10^5^ cells/cm^2^ were seeded on PCL membrane inserts in a standard 24 well cell culture plate (Sarstedt). As a control for decreased metabolic activity, cells were exposed to 20% dimethyl sulfoxide (DMSO, Merck) in cell culture media for 24 hours prior to performing the MTT assay. Then 50 μL of MTT solution (0.5 mg/mL) was added to each well, and the cells were incubated in the incubator for 1 hour. The supernatant was then aspirated, and the formazan was dissolved in 2-propanol with 0.1% Triton X-100. Absorbance was measured at 570 nm, using a plate reader (BioTek Cytation 5). Experiments were done in triplicates.

### Cell seeding on lung a chip device

MLE-12 cells were detached by trypsin, centrifuged and resuspended in cell culture media at a concentration of 3 × 10^6^ cells/mL. Prior to cell seeding, the lung on a chip device and the tubing were sterilized with 70% ethanol via flushing through the top and bottom channel inlets for 10 minutes and then rinsed 2 times with PBS. The assembled lung on a chip devices were then incubated with cell culture media for 30 minutes. After incubation, the media was removed from the channels before seeding the cells in the top channel at a cell seeding density of 3.0×10^5^ cells/cm^2^. After adhesion of cells in an incubator (37°C, 5% CO_2_) for 30 minutes, complete media was added to the bottom channel and the chips were incubated further for 72 hours until the cells formed a monolayer. The cells were then synchronized via serum starvation for 24 hours in the incubator prior to all experiments.

### Unstretched condition

FBS free culture media with 15mM HEPES (4-(2-hydroxyethyl)-1-piperazineethanesulfonic acid) (Gibco) in DMEM/F12 supplemented with 1% P/S was used for all experiments performed in the lung on a chip devices. HEPES buffer was used to ensure that the media maintains physiological pH while outside of the incubator during dynamic conditions. The chips were then placed in a dry metal bead bath (Lab Armor, M706) at 37°C and media were flowed through the bottom channel using a syringe pump with a flow rate of 7 mL/h for 2 hours while the air channel was closed. At the end of the experiments, membranes were fixed in the lung on a chip device with 10% formalin (Sigma Aldrich) and 2.5% glutaraldehyde (Sigma Aldrich, EM grade) for immunofluorescence staining and SEM analysis, respectively, and further processed as described below.

### Stretched condition

During the stretched conditions, FBS free and HEPES containing DMEM/F12 (supplemented with %1 P/S) was flowed through the bottom channel 7 mL/h as described above. The outlet of the top channel (air) was blocked and the airflow via the syringe pump was set to 0.1Hz rhythmic motion by first pumping in and out 150 µL using a flow rate of 108 mL/h for 2 hours. Membranes were immediately fixed in the lung on a chip device at the end of the experiment in 10% formalin and 2.5% glutaraldehyde.

### LDH cytotoxicity assay

Cell membrane integrity was measured via the lactate dehydrogenase activity (LDH) release assay (Roche Cytotoxicity Detection Kit). Media from the bottom channel was continuously collected in 15 minutes intervals (both for the unstretched and stretched mimicking VILI). 100 μL of media was transferred into 96 well plates and the LDH assay was performed according to the manufacturer’s protocol. In brief, the LDH reagents were mixed and added in a 1:1 (v/v) ratio with collected media and incubated for 45 minutes at room temperature. After incubation, the absorbance was measured at 490 nm using a plate reader (BioTek Cytation 5).

### SEM analysis of the membranes with cells

At the end of the experiments, membranes with cells were fixed with 2.5 wt% glutaraldehyde (Sigma Aldrich) at 4°C overnight. The membranes underwent dehydration via a graded ethanol series followed by chemical drying with hexamethyldisilazane (HMDS, Tedpella) and were sputter coated with 10 nm platinum/palladium (4/1) (Quorum Q150T ES) before being mounted and examined in a Jeol JSM-7800F FEG-SEM.

### Immunofluorescence staining

Samples from the unstretched and stretched (VILI) conditions were fixed in 10% formalin overnight at 4°C. Following cell permeabilization and blocking, the samples were incubated with YAP/TAZ (D24E4 Rabbit mAb, Cell Signaling Technology) primary antibodies (1:200) overnight at 4°C, washed three times with PBS with 0.1% Tween 20 (VWR International AB) (PBST), and then incubated for 60 minutes at room temperature with Alexa Fluor 647 goat anti-rabbit secondary antibody (1:750) (Thermo Fischer Scientific). The samples were washed three times with PBST and then incubated with 4′,6-diamidino-2-phenylindole (DAPI, 5 µg/mL in PBS) (VWR) and phalloidin (1.5 µM in DMSO) (Thermo Fischer Scientific) solution for 30 minutes. The samples were washed with PBS and imaged using a Nikon A1+ confocal system immediately. The images were analyzed with Arivis Vision4D software.

### Statistical analysis

Biological data were obtained after three independent experiments, each conducted in triplicates. Student t-tests were performed to compare the means of two normally distributed groups (stretched and unstretched). Statistics were performed and graphs were generated using GraphPad Prism 9.1.0 (GraphPad Software). P-values ≤ 0.05 are represented with (*), p-values ≤ 0.01 with (**), p-values ≤ 0.001 with (***) and p-values ≤ 0.0001 with (****).

## Results and discussion

### Fabrication of the mold, the device, and nanofibrous membrane selection

The majority of current lung on a chip devices, as well as other organs on chips, are often produced using patterned molds made from photolithography followed by PDMS casting. Photolithography offers great resolution however requires having access to dedicated and costly infrastructure such as clean-room facilities, which is not readily available for most biologically focused laboratories who are increasingly becoming the main end-users of organ-on-chips. Optimization of organ-on-a-chip devices is particularly important for maximizing their potential to reduce or replace animal experiments when primary cells are used, including those from human. 3D printed molds offer an opportunity for rapid and continuous customization for many applications and the resolution of low cost and commercially available printers has improved dramatically in the last few years, now reaching micrometer precision. Therefore, 3D printing has emerged as an alternative way to create molds for subsequent PDMS casting to generate organs on a chip, including lung. ^30^

We designed a 3D printable file in the free and open source 3D Creation Software Blender to generate molds for subsequent PDMS casting. The top and bottom part of the mold had outer dimensions of 50 mm x 40 mm and inner dimensions of 40 mm x 30 mm. The bottom part of the chip has a thickness of 1.75 mm and a top part that is 5 mm with the same outer dimensions as the original mold (**SI Figure 1**). The channels have the dimensions of 1.5 mm x 1 mm x 10 mm (**Figure 1a**).

Next, we sought to add a porous membrane which allows for air liquid interface culture and mechanical stretch to create our lung on a chip device. The majority of the previously designed lung on a chip devices have used a PDMS membrane which does not recapitulate the nanofibrous and porous extracellular matrix (ECM) found in the basement membrane and interstitial spaces of the alveolar region of the lung.

Electrospun membranes have a fibrous structure, which mimics the native lung ECM and the morphology and diameter of the fibers can be controlled via careful selection of electrospinning parameters. Importantly, the fiber diameter and density are known to affect properties such as cell attachment but also nutrient transport through the membrane.^31, 32^ Thus, we sought to explore the feasibility of using an electrospun nanofibrous PCL membrane in between the two PDMS channels to fabricate our lung on a chip device (**Figure 1b**). We used commercially available membranes provided by Cellevate AB (Sweden) with a fiber diameter distribution of 441±1490 nm (assessed via image analysis from SEM images, n=3) (**Figure 1c**). After assembling the microfluidic devices, we performed flow experiments to evaluate the robustness of the produced chips. The chips were connected to a volume-controlled syringe pump to generate perfusion of cell culture media though the bottom and top channels with a flow rate of 7 mL/h. The seal between the inlets/outlets and the PDMS was sufficient to prevent any leakage and no leakage was observed between the channels, indicating that the nanofibrous membrane could be used to form a barrier between our two channels under these flow conditions (**Figure 1b**).

### Characterization of elastic membrane properties and validation of membrane deformation

In addition to external cues from local microtopography and microenvironment, cells also sense and respond to mechanical stimuli in their environment such as mechanical stretch. Alveolar cells in the human lung are exposed to cyclic stretch between 4-12% with a frequency of 0.2 Hz during normal conditions.^1^ Under pathological conditions, alveolar distension can increase to 20% due to inhomogeneous air flow and cause injury to the lung.^33^ Additionally, mechanical ventilation (MV) is known to be a life-saving therapy for critically ill patients, where the lungs can be exposed to prolonged positive pressure in order to try and achieve sufficient oxygen delivery and gas exchange. This increase in positive pressure can result in baro-or volutrauma significant lung injury characterized by inflammation and subsequent fibrotic remodeling of the lung tissue, termed VILI.^34^ Due to the types of membranes which have been used in previous lung on a chip devices, linear strain up to only 10% has previously been possible.^15^ Therefore, we aimed to test whether our lung on a chip device could be used for studying the introduction of cyclic positive pressure on ALI cultured lung epithelial cells, with the aim of mimicking the effects of mechanical ventilation (i.e. VILI) seen in patients. First, we characterized key mechanical properties of elastic modulus and yield strain in the nanofibrous electrospun PCL membrane. Stress–strain curves performed under static loading of the electrospun PCL membranes showed that the membrane has an elastic modulus of 2.51±0.67 MPa and exhibited linear elasticity up to 14.4±4.2% yield strain **(Figure 2a)**, which is in good agreement with previously published electrospun PCL membranes of similar nanofiber morphology (∼3.0 MPa).^24^ Next, we sought to model the membrane deflection across different potential applied pressures using COMSOL simulations to achieve pathophysiological strain level. The deflection of the membrane increased with increasingly applied pressure and reached a maximal z-axis deflection of 532 μm at a positive pressure of 1.3 atm (**Figure 2b**). The deflection profile of the membrane is estimated to be parabolic and the calculated maximal strain is ∼25% which is similar to levels estimated to be present in VILI in the human lung^35^. The stretchability of the electrospun PCL membrane was tested by applying a cyclic positive pressure of 1.3 atm to the apical side of the chip. The air-filled chip was cyclically stretched while monitoring the membrane in real time, using an inverted light microscope. The membrane deflects and subsequently relaxes during injection and withdrawal of air symmetrically in opposite directions when the bottom channel is air (**SI, movie 1**) or water filled (**SI, movie 2**). Next, we experimentally determined the membrane strain and compared this with our simulation data. The displaced volume during applied strain was determined by measuring the linear displacement of a liquid plug inside tubing connected to the outlet of the bottom channel (**Figure 2c**). The displaced volume was measured to 5 μL per cycle, which corresponds to an approximate calculated vertical membrane deflection of 496.5±14.8 μm, indicating that our computational modeling and experimental data are well correlated.

**Figure 2:**
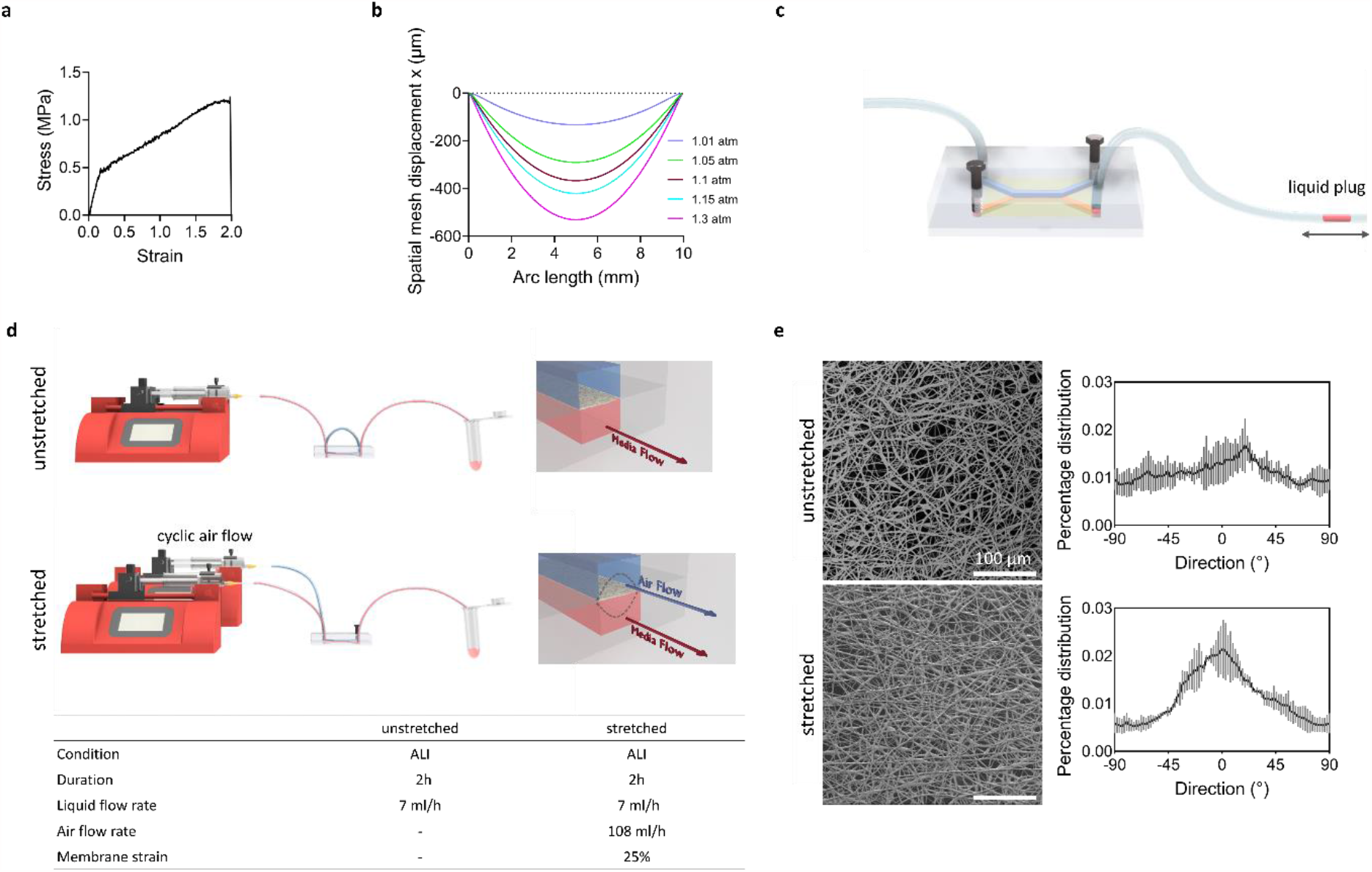
**a)** Stress-strain curve of PCL membrane. The membrane has a yield strain of 14.4±4.2%, failure strain of 173±28% and elastic modulus of 2.51±0.67 MPa (n=4). **b)** Numerical simulation of the deflection of the electrospun PCL membrane into the liquid channel as a function of applied pressure. **c)** Schematic representation of the experimental set-up for the measurement of membrane deflection. The membrane deflection was estimated based on the displacement of the liquid plug (colored water) every injection and withdrawal of the air from the top channel. **d)** Schematic of the air-liquid interface (ALI) models. Unstretched ALI model was set up by only maintaining media flow from the bottom of the channel (7 mL/h) and for stretched condition in addition to media flow air was pumped through (108 mL/h) the upper channel with a frequency of 0.1 Hz. **e)** Representative SEM images and histograms depicting fiber orientation distributions generated in ImageJ (n=3).

The main ECM constituents are collagens, elastin, and proteoglycans and each of these differentially contributes to the mechanical properties of the lung tissue giving it both elasticity and strength. Previous work has shown that collagen and elastin fiber networks deform and reorient in the direction of macroscopic strain compared to unstretched controls.^36^ As collagen is the main component giving lung tissue its mechanical strength, loss of collagen or its reorientation can cause lung injury and loss in organ function. Furthermore, collagen is known to remodel after lung injury and if it is not properly restored, including correct orientation, this can lead to the development of lung diseases.^36^ Therefore, we next sought to determine whether our electrospun PCL membranes could recapitulate this feature of the native ECM environment under conditions of stretch. We compared the orientation of PCL fibers after applying cyclic mechanical stretch at pressures yielding 25% maximal strain and compared these to unstretched controls. We found that similar to collagen and elastin fibers, irreversible fiber alignment occurred after stretching the electrospun PCL membrane with a 25% cyclic strain for 2 hours as the fibers became aligned with the applied strain (**Figure 2e**). Strain induced fiber alignment has been previously observed on electrospun PCL scaffolds.^37^ As we observed linear elasticity deformations at 14.4±4.2% yield strain in our tensile testing (**Figure 2a**), this suggests that the 25% mechanical strain we induced in our membrane is sufficient to cause plastic deformation of PCL membranes at the bulk level which is a result of fiber alignment.

### Influence of mechanical stress on distal epithelial cells

Next, we validated whether the commercially available electrospun PCL membranes can support cell adhesion, viability and proliferation of distal lung epithelial cells as this is one of the major sites of injury in VILI. PCL is a biocompatible synthetic polymer, however, it does not have molecular cues for cell binding.^1, 38^ Thus, the membranes were first incubated with complete media containing FBS which is known to contain proteins with cell binding motifs, such as fibronectin, prior to cell seeding. We found that complete media incubated PCL membranes supported the formation of a confluent monolayer of metabolically active, distal lung epithelial cells after 3 days of *in vitro* culture (**Figure 3a-e**). As a measure of cytocompatibility, the metabolic activity of the MLE-12 cells was tested with the MTT metabolic activity assay. The MTT assay showed that the cells were metabolically active compared to cells which were treated with DMSO, a substance known to inhibit metabolic activity in cells **(Figure 3f)**. This indicates that the electrospun PCL membranes do not release or leach significant amounts of cytotoxic materials (such as residual solvents or monomer) when incubated with the cell culture medium.

**Figure 3:**
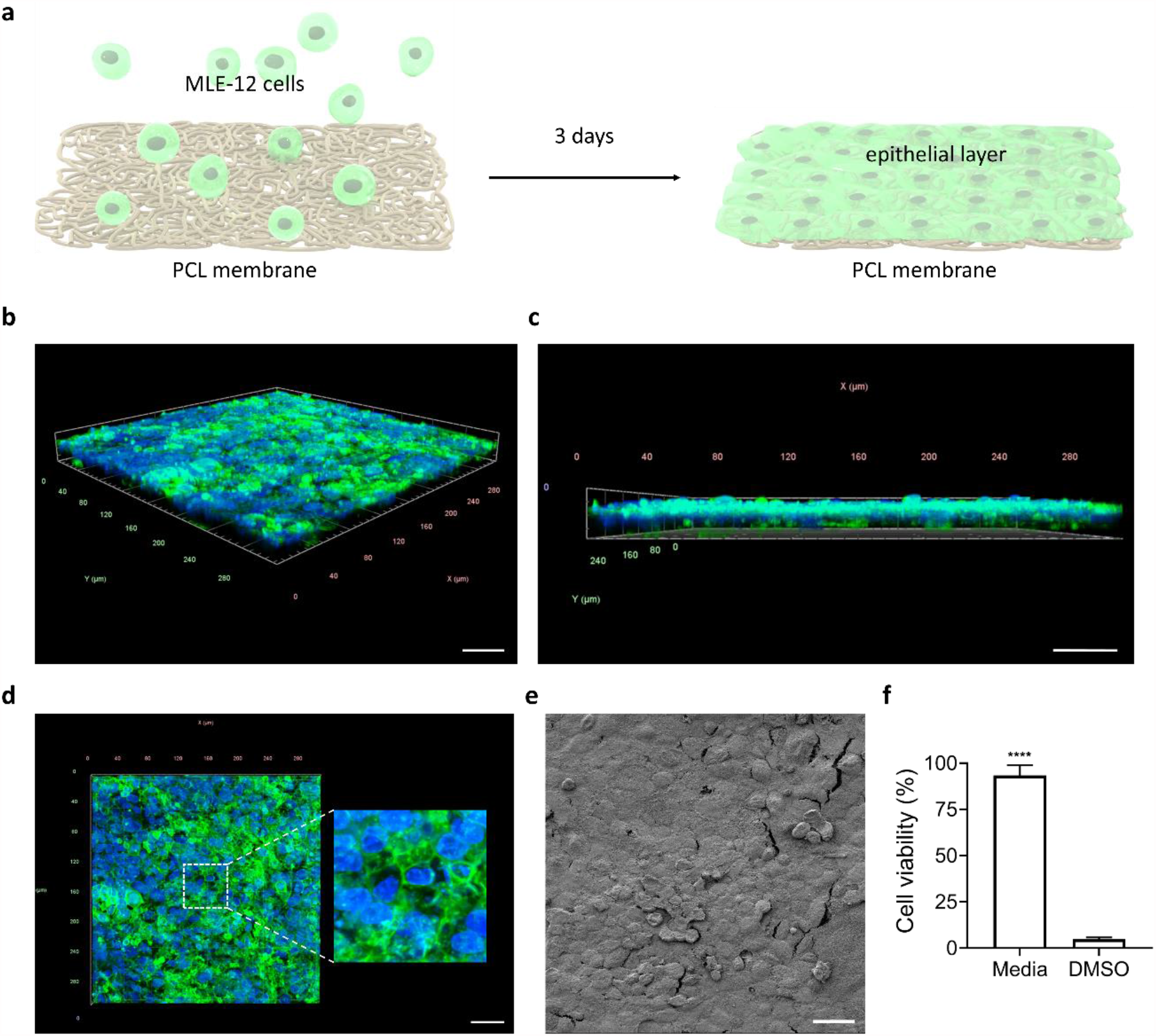
**a)** Schematic illustration of MLE-12 alveolar epithelial cell culture on electrospun PCL membrane. Monolayer of MLE-12 cells grown on PCL membrane in 3 days stained with nuclei (blue, DAPI) and actin staining (green, phalloidin) visualized by confocal microscopy; **b)** angled view **c)** front view, and **d)** top view (scale bar 40 µm). **e)** SEM image at day 3 confirms the presence of cell monolayer on the PCL membrane (scale bar 10 µm). **f)** Cell viability of the MLE-12 cells at day 3 was assessed with the MTT metabolic activity assay. As a control for decreased metabolic activity, the cells were exposed to 20% DMSO in cell culture media after 24 hours before the MTT assay. Stars show the statistical significance of change between the groups. Data are expressed as mean ± SD (n = 3) ****p < 0.0001 versus control group. Data represent three independent experiments performed with three different chips.

Next, we sought to evaluate the potential of our lung on a chip to model injurious mechanical ventilation. Under normal physiological conditions, epithelial integrity is maintained due to the force balances between the cell-cell and cell-matrix attachments. This balance is achieved through constant homeostatic modulation of cell-cell junction proteins as well as integrin and focal adhesion forces which are the point of attachment for cells on the ECM. When the adhesive forces between cells or between cells and the ECM are not sufficient to resist mechanical stretch, monolayer disruption occurs. At a tissue level, the loss of epithelial barrier integrity in the distal lung results in leakage of liquid and passage of macro-molecules and inflammatory cells between the capillary and alveolar airspace.^39^ This loss of barrier function is a hallmark of VILI.^40^ On a cellular level, cells may also be mechanically destroyed or injured during excessive shear stresses and release soluble mediators which can further drive inflammation.

Therefore, we investigated whether we could induce damage via pathophysiologic stretch of epithelial cells in our lung on a chip device. To recapitulate an epithelial barrier, distal lung epithelial cells were first seeded on the apical side of the fibrous PCL membrane in the lung on a chip device and grown to confluence under submerged conditions (**Figure 4a-d**) to build a homogeneous monolayer just prior to generating ALI conditions by removing media on the top channel and filling with air. Cells were exposed to a shear stress of 54 dynes/cm^2^ on the basal side of the membrane using a fluid (media) flow rate of 7 mL/h for 2 hours. In conditions aiming to replicate mechanical ventilation induced lung injury, the cells were exposed to 25% cyclic strain at 0.1Hz for 2 hours by over pressurizing the top channel with air, while the media was flowing through the bottom channel (**Figure 4e**).

**Figure 4:**
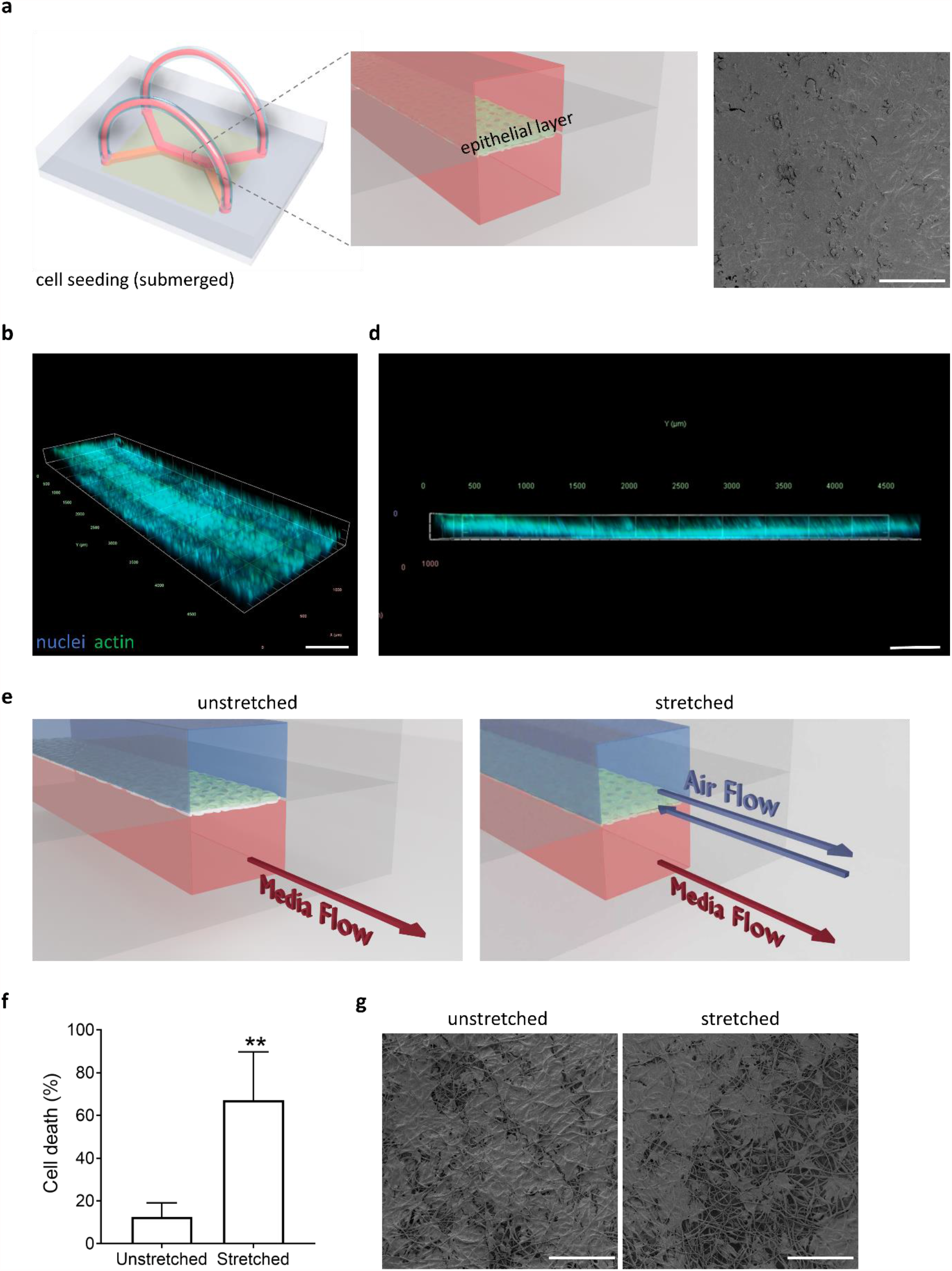
**a)** Monolayer growth of MLE-12 cells in the lung-on-a-chip device. **b)** Scanning electron image showing the presence of a confluent lung epithelial layer on the membrane after 3 days under static submerged cell culture conditions. The nucleus and actin filaments of the cells were stained using DAPI (blue) and phalloidin (green), respectively. **(c-d)** Confocal fluorescence microscopy images of the angled and the left views of the membrane in between of the channels confirms the presence of MLE-12 monolayer on the electrospun PCL membrane. **e)** After the formation of confluent MLE-12 layer, the cells were cultured in ALI conditions. Chips were maintained under constant flow of fresh medium (7mL/h) for both unstretched and stretched condition for 2 hours. Membrane stretch was achieved by blocking the outlet of the air channel and flowing air (108 mL/h) with a frequency of 0.1 Hz for 2 hours (stretched condition). **f)** Cell death of the MLE-12 cells was assessed with LDH cytotoxicity assay. **g)** SEM images show changes in the epithelial layer after the cell layer was exposed to constant flow of media (7mL/h) (unstretched) and also cyclic membrane deformations (25% strain, 0.1 Hz) (stretched) for 2 hours. Data are expressed as mean ± SD (n = 3) **p < 0.0043 versus unstretched group. Data represent three independent experiments performed with at least three different chips. Scale bars: 100 µm.

Similar to previous work^7, 19, 41^, we found that pathophysiologic stretch caused an increase in the concentration of the cell injury marker, lactase dehydrogenase (LDH), in the basal media, indicating that our pathophysiologic stretch conditions induced cellular injury in the lung on a chip device and that this could be measured in the media containing channel, similar to LDH measurements in the blood of animals and patients with VILI (**Figure 4f**).^42^ Furthermore, we found that our pathophysiologic stretch conditions caused disruption of the epithelial monolayer, as evident by a pronounced loss in cellular confluence following pathophysiologic stretch as compared to the unstretched condition (**Figure 4g**). On the other hand, the majority of the monolayer remained intact under ALI conditions and basal media perfusion in the unstretched condition; however, some cell loss can be observed from the membrane surface, likely due to media flow from the bottom channel.

Having established that our model recapitulated some of the main features of mechanical stretch induced injury (i.e. monolayer loss and increased concentrations of LDH), we next sought to validate if our lung on a chip could be used to explore cellular pathways known to be altered with mechanical stretch. Alveolar cells are known to become damaged from high levels of strain and alveolar injury is increasingly recognized as a major drive of disease pathogenesis.^43^ Cells are known to be capable of transducing mechanical changes in their cellular environment into biochemical signals through so called mechanotransduction pathways. These pathways allow cells to retain tissue level homeostasis and for cells to adapt to their environment but the same pathways can become activated in pathologic conditions and be further drivers of disease themselves. One such pathway which has received increased attention is through the mechanotransducers YAP/TAZ which have been shown to be responsive due to changes in substrate stiffness, cell-cell junctions as well as mechanical stretch.^39^ Furthermore, YAP/TAZ are increasingly recognized as playing a central role in alveolar epithelial cell behavior during development, normal postnatal regeneration, and can also be pathologically activated after lung injury.^44^

Therefore, we examined whether pathophysiologic stretch in our chip can induce YAP/TAZ translocation from the cytoplasm to the nucleus. We found that YAP/TAZ translocated to nuclei when the epithelial cells were exposed to cyclic pathological strain, while YAP/TAZ remained mainly in the cytoplasm in the unstretched conditions (**Figure 5**). Thus, pathophysiologic stretch of cells seeded on a nanofibrous membrane in our lung on a chip is able to induce cellular level changes in known mechanotransducers and establishes the feasibility of using this lung-on-a-chip device to monitor cellular level changes.

**Figure 5:**
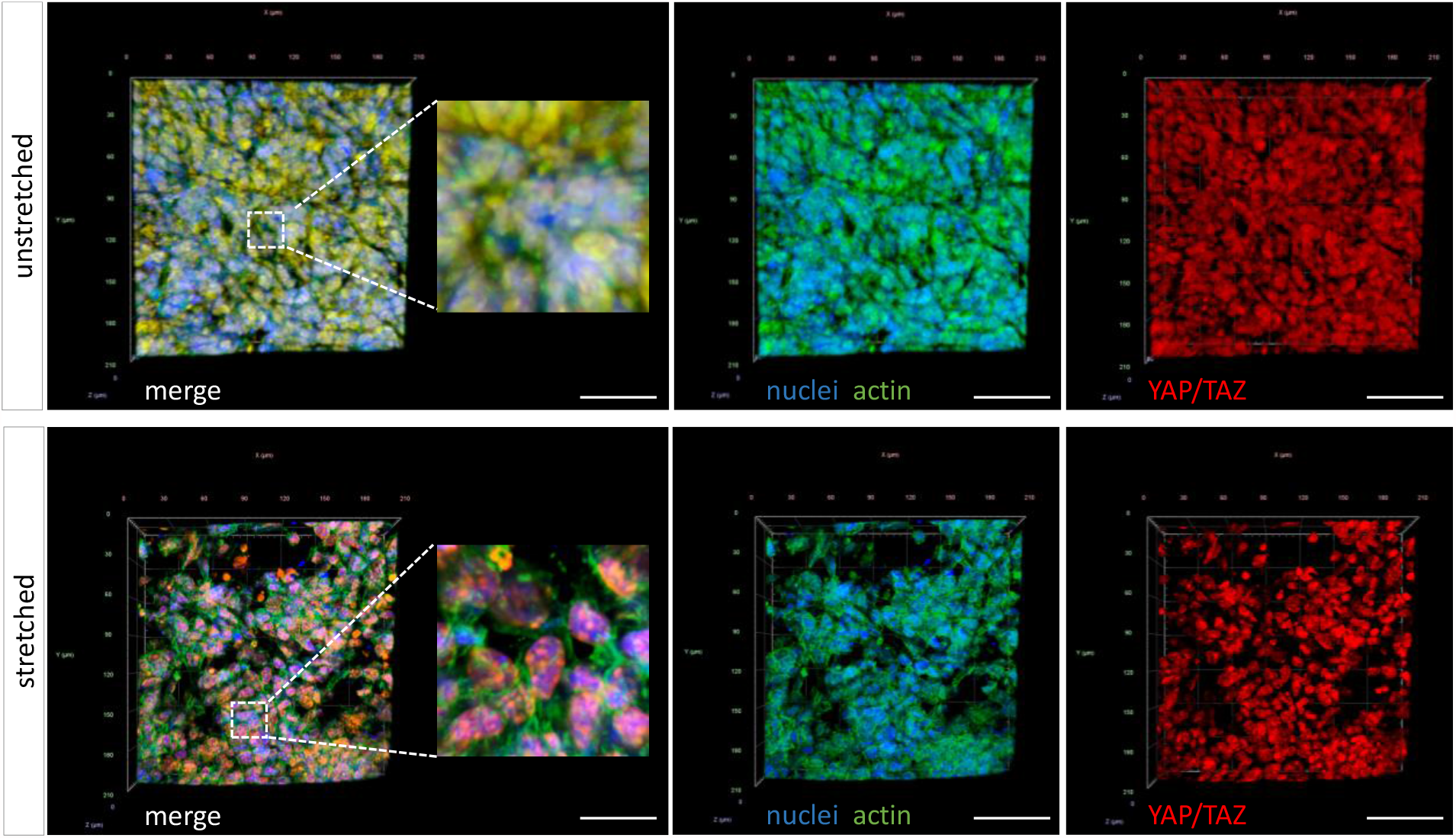
Effect of stretching of MLE-12 cells on YAP/TAZ localization. Cells seeded on the lung-on-a-chip device and grown to monolayer for three days. Serum starved cells were subjected to unstretched and stretched conditions, fixed and then stained with DAPI (nuclei, blue), phalloidin (actin, green) and YAP/TAZ antibody (red) for immunofluorescence imaging (scale bar 60 µm). YAP/TAZ was widely visible in the cytoplasm under unstretched condition (appears yellow in the merged panel due to colocalization of F-actin (green) and YAP/TAZ (red)). The cells displayed preferential nuclear YAP/TAZ localization (clear nuclear staining, including punctae) by applying pathophysiological cyclic stretch. Data represent three independent experiments performed with three different chips.

## Conclusion

In this study, we demonstrated the potential of 3D-printed molds to fabricate a simple and cost-effective lung on a chip device that uses a commercially available, stretchable and nanofibrous membrane to mimic VILI. The device can be manufactured in any lab that has access to a 3D printer and means to perform soft lithography, making it easily available to anyone. 3D printing enables fabrication of devices in a short period of time (from days to a few hours) and is amenable to tailorability in terms of device design. Previous *in vitro* cell culture experiments with alveolar epithelial cells cultured under physiologic ALI conditions have utilized non-fibrous stretchable PDMS membranes or fibrous membranes made of natural polymers such as collagen and have shown an upper limit of around 10% mechanical strain, making studies of VILI challenging.^12, 15^ Here we used a thin electrospun nanofibrous PCL membrane to mimic the natural basement membrane. We found that this nanofibrous PCL membrane supported the growth of monolayers of lung epithelial cells and had properties that enabled us to apply pathophysiological stretch to mimic VILI *in vitro*, including mimicking of collagen rearrangement at higher mechanical strains. Finally, we were able to confirm that pathophysiologic stretch induces cell injury and subsequently cell death compared to unstretched controls. In addition to cell death, cyclic pathophysiologic stretch induced cellular level changes in known mechanotransducers of YAP/TAZ. As an outlook, the lung on a chip device introduced here should be tested with primary human alveolar epithelial cells, human endothelial cells and macrophages to create alveolar-capillary barrier models in-vitro, including for personalized medicine approaches.^45-47^ Furthermore, it will be interesting to evaluate the potential of our lung on a chip manufacturing pipeline, including this chip geometry, to model environmental stimuli in the context of mechanical stretch with cigarette smoke, e-cigarettes, pathogens and particle exposure mimicking pollution and occupational exposure. The combined advances we demonstrate with this highly adaptable and low-cost lung on a chip fabrication technique could pave the way for a significantly improved platform to study diverse behavior under mechanical ventilation as well as to evaluate drug responses to prevent injury during the mechanical ventilation.

## Acknowledgement

The authors thank Cellevate AB, Sweden for providing nanofibrous PCL membranes and helpful discussions throughout the project. We also thank Sebastian Wasserstrom from LBIC for technical assistance with the SEM and confocal microscopy. We also would like to thank all the Lung Bioengineering and Regeneration group members for helpful discussions. This project has received funding from the European Research Council (ERC) (grant agreement No 805361) (D.E.W.). Further support is acknowledged from the Knut and Alice Wallenberg foundation (D.E.W.). S.T. is supported by a RESPIRE3 Postdoctoral Fellowship supported by the European Respiratory Society and the European Union’s H2020 research and innovation programme under the Marie Sklodowska-Curie grant agreement (Grant No. 713406). JPB was supported by a grant from the Swedish Research Council (grant number 2019-02355).

## Supporting Information

**Figure S1.**
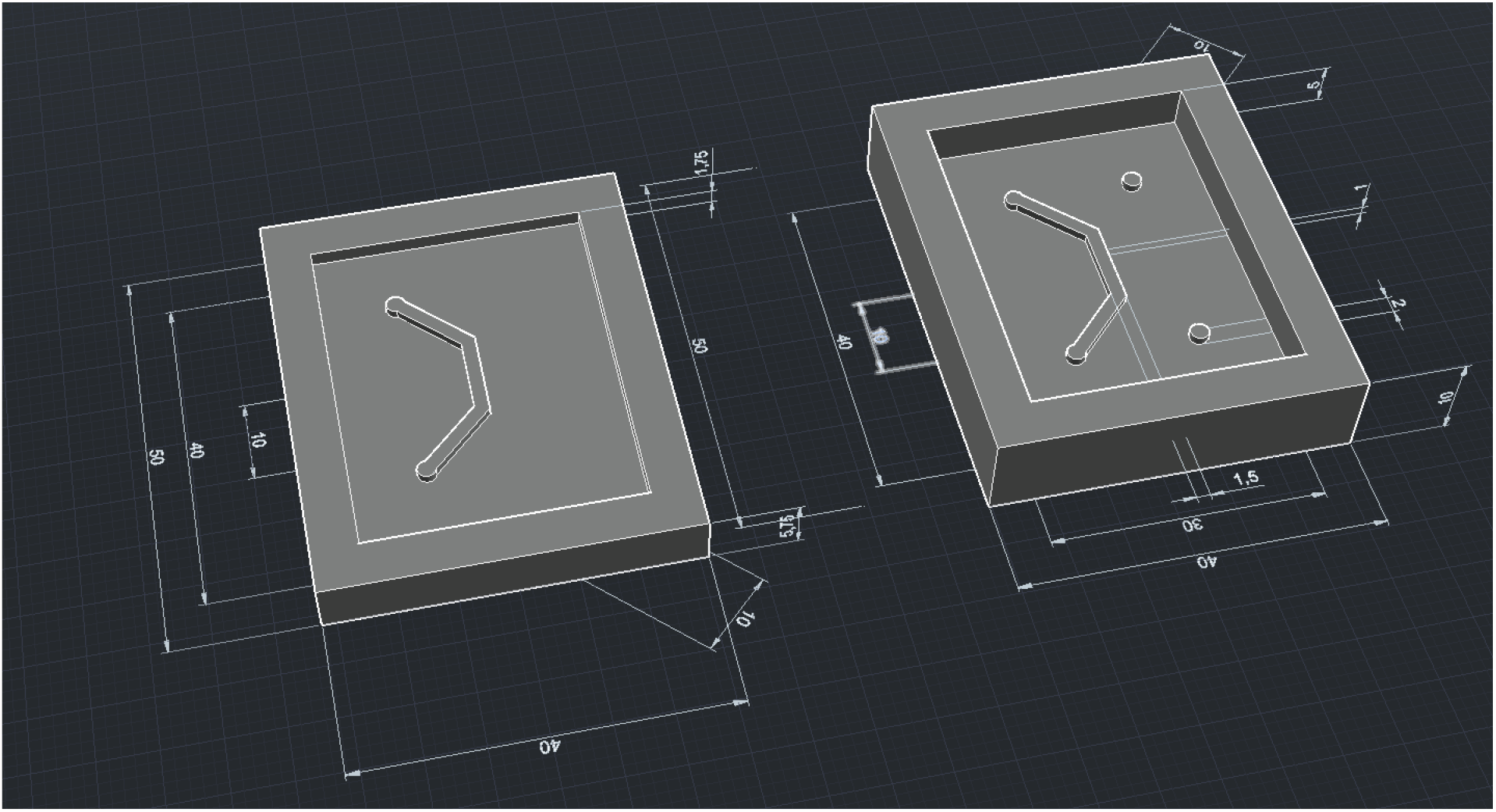
CAD drawings of the 3D printed mold (units: mm)^1^

**Movie S1**. Nanofibrous electrospun PCL membrane deflection during injection and withdrawal of air in a fabricated lung on a chip device without fluid in the lower channel.

(Viewable: https://youtu.be/6NKYy4lxNCc)

**Movie S2**. Nanofibrous electrospun PCL membrane deflection during injection and withdrawal of air in a fabricated lung on a chip device with fluid(water) in the lower channel.

(Viewable: https://youtu.be/gBlO3jGupAo)

## Notes

### Competing Interest Statement

The authors have declared no competing interest.

